# Challenging the Raunkiaeran shortfall and the consequences of using imputed databases

**DOI:** 10.1101/081778

**Authors:** Lucas Jardim, Luis Mauricio Bini, José Alexandre Felizola Diniz-Filho, Fabricio Villalobos

## Abstract

1. Given the prevalence of missing data on species’ traits – Raunkiaeran shorfall — and its importance for theoretical and empirical investigations, several methods have been proposed to fill sparse databases. Despite its advantages, imputation of missing data can introduce biases. Here, we evaluate the bias in descriptive statistics, model parameters, and phylogenetic signal estimation from imputed databases under different missing and imputing scenarios.
2. We simulated coalescent phylogenies and traits under Brownian Motion and different Ornstein-Uhlenbeck evolutionary models. Missing values were created using three scenarios: missing completely at random, missing at random but phylogenetically structured and missing at random but correlated with some other variable. We considered four methods for handling missing data: delete missing values, imputation based on observed mean trait value, Phylogenetic Eigenvectors Maps and Multiple Imputation by Chained Equations. Finally, we assessed estimation errors of descriptive statistics (mean, variance), regression coefficient, Moran’s correlogram and Blomberg’s K of imputed traits.
3. We found that percentage of missing data, missing mechanisms, Ornstein-Uhlenbeck strength and handling methods were important to define estimation errors. When data were missing completely at random, descriptive statistics were well estimated but Moran’s correlogram and Blomberg’s K were not well estimated, depending on handling methods. We also found that handling methods performed worse when data were missing at random, but phylogenetically structured. In this case adding phylogenetic information provided better estimates. Although the error caused by imputation was correlated with estimation errors, we found that such relationship is not linear with estimation errors getting larger as the imputation error increases.
4. Imputed trait databases could bias ecological and evolutionary analyses. We advise researchers to share their raw data along with their imputed database, flagging imputed data and providing information on the imputation process. Thus, users can and should consider the pattern of missing data and then look for the best method to overcome this problem. In addition, we suggest the development of phylogenetic methods that consider imputation uncertainty, phylogenetic autocorrelation and preserve the level of phylogenetic signal of the original data.

## Introduction

Missing data are a ubiquitous feature of real-world datasets (Nakagawa & Freckleton 2008). Lack of information may limit the application of statistical analysis and can lead to biased estimates and conclusions on the phenomena of interest. In 1976, Donald B. Rubin proposed a missing-data theory to allow analysis of incomplete datasets (Rubin 1976), explaining how unbiased parameters could be estimated with missing data by considering the mechanisms causing missing data. These mechanisms were classified in three categories: missing completely at random (MCAR), missing at random (MAR) and missing not at random (MNAR). They mean, respectively, that missing values are equally probable across a dataset, probability of missing data is correlated with other variables rather than to the variable with missing data (target variable), and probability of missing data is itself correlated to the target variable and dependent on the missing data (Rubin 1976; Nakagawa & Freckleton 2008; Enders 2010; van Buuren 2012) (Fig.1).

**Figure 1.**
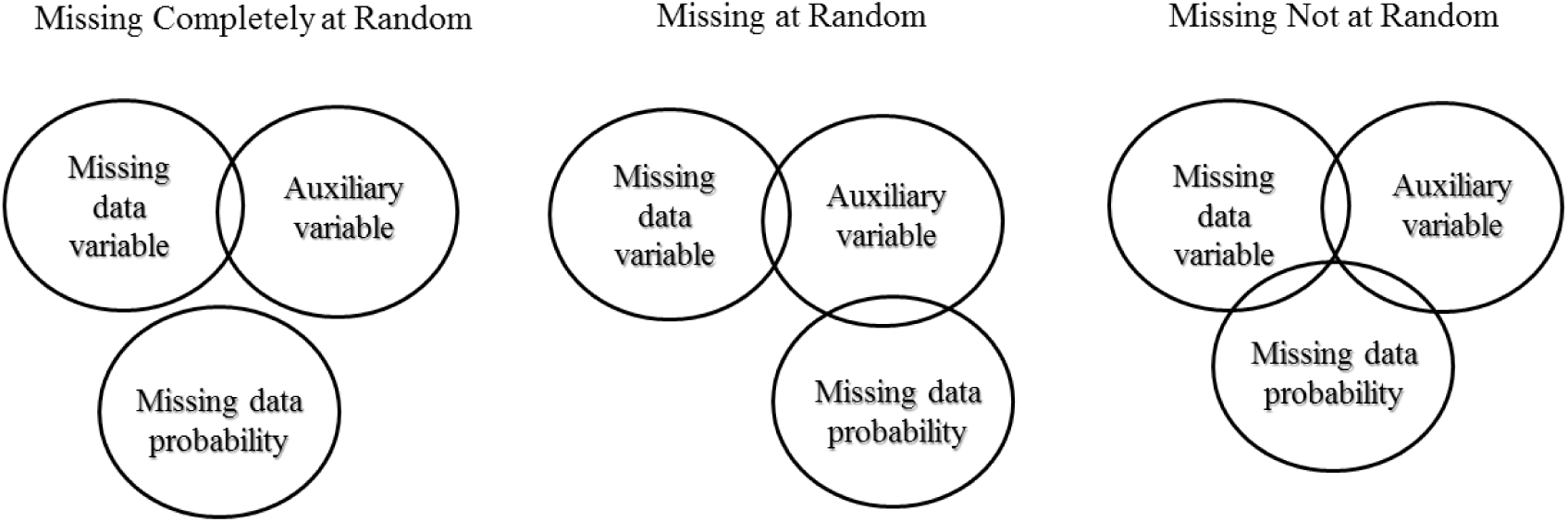
Correlation structure among variables in each missing data mechanism. Circles represent model components and their intersection represents correlation among them.

When dealing with missing data, the above mechanisms need to be taken into account before analysis (Rubin 1976). This is because different methods that handle missing data assume different mechanisms, so using them indiscriminately may bias parameters estimates (Rubin 1976; Enders 2010; van Buuren 2012). Multiple Imputation and Full Information Maximum Likelihood methods are currently regarded as the most appropriate methods to handle missing data, because they work under MAR and MCAR scenarios and provide unbiased estimates (Enders 2010). In contrast, it is very difficult to model missing data under a MNAR scenario. This is so due to the need of considering a model that represents the probability of missing values to occur and because the shape of the probability density function is not known (Enders 2010; van Buuren 2012).

Research in ecology and evolutionary biology usually requires data about species and their traits to answer different questions from community assembly and ecogeographical rules to correlated evolution, diversification rates and extinction probability, among others (Purvis et al. 2000; Webb et al. 2002; Gaston et al. 2008; Goldberg et al. 2010; Lukas & Clutton-Brock 2013; Jetz & Freckleton 2015). Thus, to facilitate research and make it reproducible and data more accessible (Reichman et al. 2011), ecologists and evolutionary biologists usually create databases that include information on huge amounts of species and their traits (e.g., (Jones *et al.* 2009; Kattge *et al.* 2011; Wilman *et al.* 2014). However, as databases become larger, the probability of having all the necessary data for all species rapidly decreases. This lack of knowledge about species’ traits and their ecological functions was recently defined as the Raunkiaeran shortfall (Hortal et al. 2015) or Eltonian shortfall (Rosado *et al.* 2015).

Owing to the ubiquity of the Raunkiaeran shortfall, some researchers are interested in filling such gaps in their databases for their own analyses but also to make them available for other researchers (Swenson 2014; Schrodt *et al.* 2015). To do so, recent studies suggest the use of phylogenetic information in the imputation process (Guénard *et al.* 2013; Swenson 2014; Schrodt *et al.* 2015). Phylogenetic information is important in imputation because closely related species resemble, on average, each other more than distantly related species. Such phenomenon is commonly known as phylogenetic signal (Blomberg *et al.* 2003). Consequently, knowing the phylogenetic position of species could, in principle, be used to perform a good estimation of missing trait values. However, the relationship between trait divergence and phylogenetic distance may be more complex (due to distinct evolutionary models) than usually assumed (Hansen & Martins 1996; Münkemüller *et al.* 2012). For instance, under Ornstein-Uhlenbeck evolutionary model traits may evolve under selection restrictions where species track a trait optimum, causing phenotypic resemblance even among phylogenetically distant species (Hansen & Martins 1996). Alternatively, under Early-burst model traits may show evolutionary rates early in species history and later the rates slow down, resulting in phylogenetically closely related species having different trait values (Blomberg *et al.* 2003; Harmon *et al.* 2010). Finally, trait evolution may happen under a drift process (e.g., Brownian motion) where species trait differences are directly correlated with time since divergence (Felsenstein 1985; Hansen & Martins 1996; Freckleton *et al.* 2002). Therefore, imputation methods should explicitly consider or assume a trait evolutionary model determining the relationship between species resemblance and phylogenetic proximity (Guénard *et al.* 2013).

Nowadays, large, imputed databases already exist that used taxonomic, ecological or allometric relationships to fill in missing values (Jones *et al.* 2009; Wilman *et al.* 2014). This highlights the need to critically evaluate the use of imputed databases given that the reliability of statistical analysis under missing data is dependent on how much values were missing in the original data, what mechanism caused data to be missing and which methods were used in the imputation process (Schafer & Graham 2002; Enders 2010; van Buuren 2012). Moreover, other problems can also arise when testing for phylogenetic signal (Cavender-Bares *et al.* 2009; Münkemüller *et al.* 2012). In such cases, if analysis were to be conducted on phylogenetically imputed data, results could be misleading given that missing values would have been already filled based on their phylogenetic structure, thus potentially inflating the level of phylogenetic signal. This potential issue can have important consequences for studies evaluating, for example, niche conservatism, trait lability, community assembly and diversification (Blomberg *et al.* 2003; Wiens & Graham 2005; Cavender-Bares *et al.* 2009; Goldberg *et al.* 2010).

Considering the current need for complete databases and the use of imputation methods to accomplish this, we evaluate how the estimation of descriptive statistics, regression coefficients and phylogenetic signal can be misled by the percentage of missing data, the particular mechanism of missing data, the model of trait evolution and the choice of methods used to handle missing values. To accommodate all of these scenarios, we use simulated phylogenies and traits under different combinations of such conditions. In addition, to address imputation accuracy, we evaluated the relationship between error caused by imputation and statistical estimation errors.

## Methods

### Phylogeny simulation

To evaluate the effect of imputing missing values into sparse databases (i.e. with missing data), we first simulated 100 coalescent phylogenies with 200 species using the function *rcoal* from the R package *ape* (Paradis *et al.* 2004). We focused on this phylogeny size because it has been considered appropriate to evaluate power and accuracy of phylogenetic analysis (Davis *et al.* 2013; Cooper *et al.* 2015), and it represents a conservative approximation to database size (e.g. several hundreds to thousands of species).

### Trait simulation

For each phylogeny, we simulated two traits: a target trait and an auxiliary trait. The first trait represented the one that would be imputed (i.e. missing-value trait), whereas the second trait represented an auxiliary trait that would be used to impute values for the target trait.

The target trait was simulated using the *rTraitCont* function from the *ape* package (Paradis *et al.* 2004). We modeled this trait under a Ornstein-Uhlenbeck evolutionary process (OU) (Gillespie 1996), because it allowed us to simulate trait evolution within a continuum from evolutionary drift (i.e. Brownian motion) to weak and strong levels of selection strength on trait evolution (Hansen & Martins 1996; Hansen 1997). Thus, we could evaluate the performance of imputation methods under different levels of phylogenetic signal. We fixed the target trait’s optimum (ϴ) to zero and the trait interspecific variation (σ) equal to one. Also, we simulated different selection strengths by varying α (selective strength) from 0 to 2, in 0.5 steps (0, 0.5, 1, 1.5 and 2). Such values covered evolutionary scenarios from Brownian motion (OU α = 0) to strong selective strength (OU α = 2).

The auxiliary trait represented a variable used to impute values into the target trait. We simulated auxiliary traits in two ways: (i) correlated with the phylogeny and (ii) correlated with the target trait but uncorrelated with phylogeny. For (i), we simulated the trait following Liam Revell (pers. comm.):

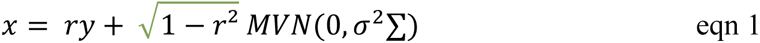

where *y* is the target trait, *x* the auxiliary trait, and *r* the correlation coefficient between both traits. We set *r* equal to 0.6 and 0.9 to explore the sensibility of our results to the strength of trait correlation. **∑** is the species covariance matrix (Felsenstein 1985; Revell *et al.* 2008) and σ² the target trait variation rate calculated as the mean of squared phylogenetic independent contrasts (Freckleton & Jetz 2009), which was estimated using the *pic* function from *ape* (Paradis *et al.* 2004). MVN means Multivariate Normal Distribution and it was simulated using the *fastBM* function from the *phytools* R package (Revell 2012). This auxiliary trait was later used when simulating the MCAR (Missing Completely at Random) and MAR.PHYLO (Missing at Random correlated with phylogeny) (see below).

For the second scenario, where the auxiliary trait is correlated with the target trait but uncorrelated with phylogeny, the auxiliary trait was simulated using equation 1 with ∑ having off-diagonal entries equal to zero (i.e. no covariance among species) and diagonal entries representing, for each species, the sum of all branch lengths from the root to the tip. We simulated MVN using the *mvrnorm* function in the R package MASS (Venables & Ripley 2002). When using this auxiliary trait to impute target trait values, we expected that using the phylogeny into the imputation methods would not improve our analysis (i.e. provide no information on missing data) since the probability of missing values would only be correlated with the auxiliary trait and not with the phylogeny.

### Missing data scenarios

To create missing data, we used the target trait simulated above and deleted different percentages of its values following three scenarios of missing data: Missing Completely at Random (MCAR), Missing at Random but phylogenetically structured (MAR.PHYLO), and Missing at Random but correlated with another phylogenetically unstructured trait (MAR.TRAIT). We created the MCAR scenario by randomly sampling a percentage (see below) of species along each phylogeny and replacing their trait values with missing values. For the MAR.PHYLO scenario, we sampled a species in each phylogeny and selected a percentage of its closest species to replace their trait values with missing values, allowing a strong missing data pattern that was phylogenetically structured. For the last scenario, MAR.TRAIT, we used the auxiliary trait (see above) to replace values in the target trait. We ordered the values of the auxiliary trait in ascending order and replaced the first percentage of values of the target trait with missing values. This represented a missing data pattern correlated with another trait, different to the target one. For each scenario, we simulated different percentages of missing values in the target trait: 5, 10, 20, 50, 70 and 90% of missing data. These percentages were chosen to represent common proportions of missing data present in highly used databases such as PanTHERIA (Jones et al., 2009) and EltonTraits (Wilman *et al.* 2014) (Fig. S1, Appendix S2).

### Imputation methods

We evaluated four methods often applied by researchers to handle missing data: imputation based on averaging values (MEAN), no imputation and simply deleting missing values (LISTWISE), phylogenetic eigenvector maps (PEM), and multiple imputation by chained equations (MICE).

We used the MEAN method to impute missing values by filling them with the average of the observed values of the target trait. Under the LISTWISE method, we did not impute values but simply deleted those species with missing values in the phylogenies before the analyses. The PEM method uses both phylogenetic eigenvectors (Diniz-Filho *et al.* 1998) and traits to impute data considering different OU processes (Guénard *et al.* 2013). We applied this method in two ways: first, using only the phylogenetic eigenvectors (PEM.notrait) and, second, using these eigenvectors and the auxiliary trait (PEM.trait). By applying the PEM method in these two ways allowed us to evaluate whether phylogenetic information alone could impute data well or auxiliary traits were necessary. Eigenvector selection and fitting of trait evolutionary models were performed using the *MPSEM* R package (Guénard *et al.* 2013) using forward selection based on the second-order Akaike Information Criterion. The MICE method simulates several possible values for missing data from a posterior predictive distribution, then runs analysis and pools results over all simulated data (van Buuren *et al.* 2006). We chose this method because it is flexible and allows imputing categorical, continuous, and non-normally distributed data (van Buuren *et al.* 2006). We applied MICE by creating 10 datasets to run our analysis over them and pooled the results. The quantity of datasets created by MICE is dependent on the percentage of missing data and more datasets can provide higher accuracy and power in the analyses (Graham *et al.* 2007; Enders 2010; van Buuren 2012). However, because our objective was simply to estimate statistical bias instead of inference power, 10 datasets can be considered appropriate (Graham *et al.* 2007). As with the PEM method, we applied MICE in two ways: only considering the auxiliary trait (MICE) and using this trait plus the phylogenetic eigenvectors selected as in PEM (MICE.phylo). We imputed data with MICE using the *mice* R package (van Buuren & Groothuis-Oudshoorn 2011).

We simulated 540 scenarios representing each combination of missing data percentage, mechanism, OU selection strength, and imputation methods. For each scenario, we simulated 100 replicates, thus producing 54000 independent results. Finally, we averaged 10 replicates for each scenario and ended up with 5400 simulations to analyze.

### Estimating Phylogenetic Signal

We calculated the phylogenetic signal (PS) in our simulated phylogenies using two metrics: Blomberg’s K (Blomberg *et al.* 2003) calculated with the *phylosig* function of *phytools* (Revell 2012) and Moran’s *I* correlograms (Gittleman & Kot 1990; Diniz-Filho 2001). For these correlograms, we created a phylogenetic distance matrix per phylogeny using the *cophenetic* function of *ape* (Paradis *et al.* 2004) and built the correlograms with the *lets.correl* function of the *letsR* R package (Vilela & Villalobos 2015). Then, based on the correlogram, we used the intercept of the following linear model as indicative of PS:

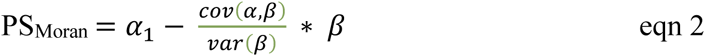

where *cov* is the covariance between the mean within-class distance and Moran’s Index, *var* is the variance of the mean within-class distance, α is the value in each correlogram distance class, and α 1 is the value in the first distance class.

### Imputation effects on phylogenetic signal

To evaluate the effect of using imputed trait data for estimating phylogenetic scenario. In particular, we estimated PS for the original target trait values before they were deleted by the missing data mechanisms and estimated PS again after they were filled by the imputation methods. Such PS delta was defined as:

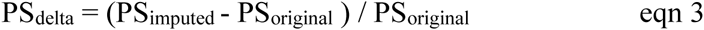

where PS_imputed_ is the PS calculated after imputing missing data, PS_original_ is the observed PS. If PS_delta_ is positive, there is a gain in PS (i.e. more PS than the original data), meaning that imputing data increased the phylogenetic structure of the target trait. Conversely, if PS_delta_ is negative, there is a decrease in PS (i.e. less PS than the original data), meaning that imputing data decreased phylogenetic structure and made species’ trait values seem more phylogenetically independent than they originally were. PS_delta_ equal to zero represents no change in trait phylogenetic structure (i.e. no imputation effect).

### Imputation effects on descriptive statistics

Traditionally, performance evaluation of imputation methods have focused on common descriptive statistics such as (mean, variance, regression coefficient) (Collins *et al.* 2001; van Buuren *et al.* 2006; Penone *et al.* 2014) instead of phylogenetic patterns. Therefore, we also evaluated the effect of imputed data on the estimation of such descriptive statistics. We calculated the mean and variance of the target trait as well as the regression coefficient (Ordinary Least Square) between the target trait and the auxiliary trait, before producing missing data and after imputing such data. Next, we measured the estimation error for these statistics as the mean squared error (MSE), as below:

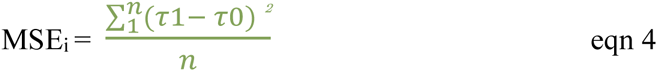

where τ1 represents the statistics calculated over imputed traits, τ0 is the statistics calculated from original traits, *n* means the number of simulations averaged to result ith MSE value.

### Imputation error

To measure the potential error introduced by imputation methods, that is the deviation between imputed and original data, we followed Penone *et al.* (2014) and used the normalized root mean squared error (NRMSE):

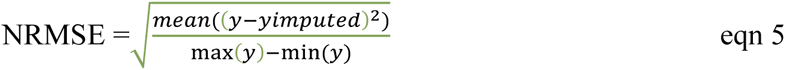

where *y* is the original trait value, *y*_imputed_ is the imputed value, max(*y*) and min(*y*) are the maximum and minimum values of the original trait, respectively. NRMSE varies between 0, no estimation error, and 1, maximum error (Oba *et al.* 2003).

### Overall analyses

We were also interested on evaluating the effects of percentage of data missing, missing data mechanism, OU selection strength, and imputation methods as factors influencing the abovementioned effects of imputation (estimation errors: PS_delta_ and MSE of descriptive statistics). To do so, we built linear models with these factors (e.g. percentage of data missing) and their interactions as predictors and estimation errors, separately, as individual responses. We specified the models using the *dredge* function from the *MuMIn* R package (Bartón 2016) and ranked the different models using delta AICc (Burnham & Anderson 2002). In addition, given concerns on the accuracy of imputation methods (Guénard *et al.* 2013; Penone *et al.* 2014), we also evaluated the relationship between imputation error (NRSME) and estimation errors (PS_delta_ and MSE) caused by imputation. All simulations and analysis were run in R 3.2.2 (R Core Team 2015).

## Results

In our simulations we found that differences in estimation errors were dependent on missingness mechanism, imputation method, evolutionary model, percentage of missing data and statistics being estimated. Moreover, imputation errors showed different results between trait correlations (target vs. auxiliary trait; *r*) of 0.6 and 0.9, but descriptive statistics and phylogenetic signal errors did not show different results concerning this correlation. Therefore, we present all results for *r* = 0.6 and only those for *r* = 0.9 corresponding to *Imputation error* (see below). Full results for *r =* 0.9 can be found in the Appendix S1.

Not surprisingly, our results showed a clear tendency of increasing error in estimating phylogenetic signal and descriptive statistics as the percentage of missing data gets larger (Fig. 2). We did not identify a clear threshold in the amount of missing data that would guarantee lower statistical errors. For the best imputation methods (MICE.phy, PEM.trait, PEM.notrait; see below) lower errors were possible for as low as 30% and up to 70% of missing data in the target trait.

**Figure 2.**
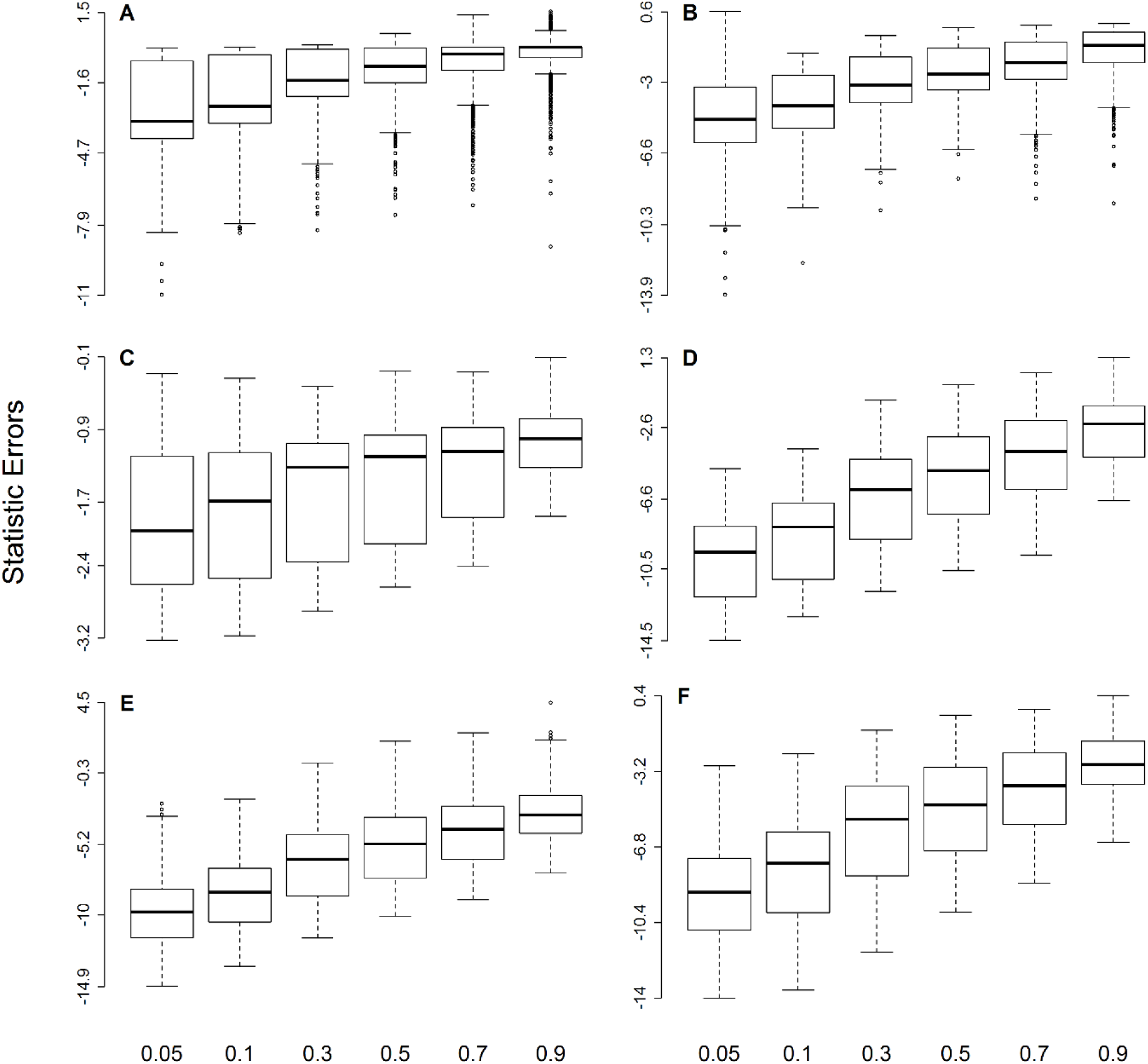
Missing data percentage and statistic estimation errors. (A) Logarithm of absolute average proportions of Blomberg’s K change after imputation or deletion, (B) Logarithm of average proportion of Moran’s Correlogram values change, (C) Imputation error measured as logarithm of average NRMSE, (D) Logarithm of MSE of trait mean, (E) Logarithm of MSE of trait variance and (F) Logarithm of MSE of regression coefficient.

When data were missing completely at random (MCAR), most imputation methods showed good performance (Fig. 3-5; Fig. S2 and S3, in Appendix S2), except the MEAN method. Nevertheless, when estimating Blomberg’s K only LISTWISE and PEM.trait resulted in low proportional changes (Fig. 3). For mean, variance and regression coefficient MSE, imputation methods worked better when data were missing at random but correlated with another trait (MAR.TRAIT) than when data were missing and phylogenetically structured (MAR.PHYLO) (Fig. 5; Fig. S2 and S3, Appendix S2). Nevertheless, the lowest proportional changes in Blomberg’s K and PS_Moran_ were observed under the MAR.PHYLO scenario (Fig. 3 and 4).

**Figure 3.**
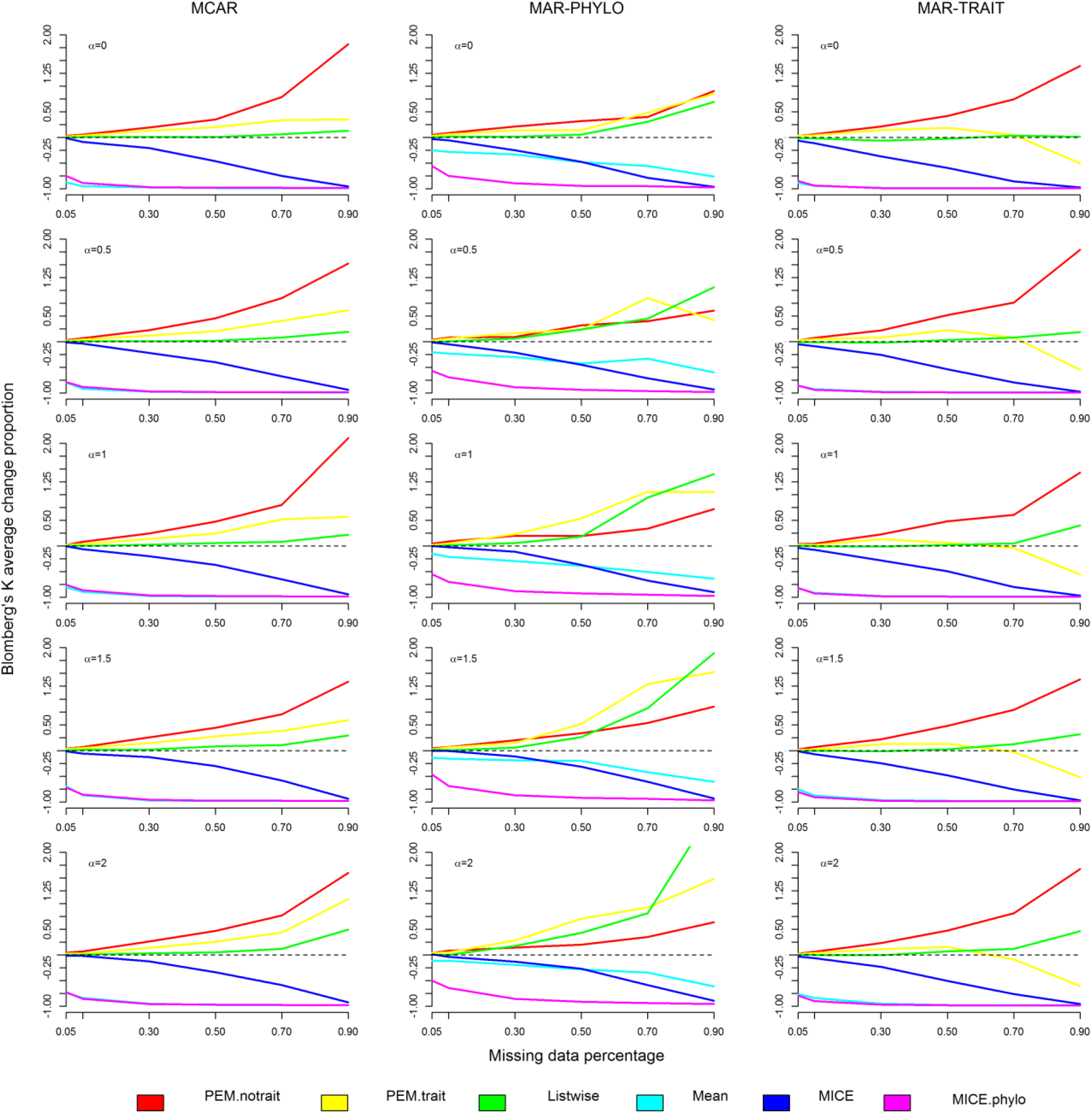
Blomberg’s K average change proportion under different methods, OU selective strength, missing data percentage and mechanisms.

**Figure 4.**
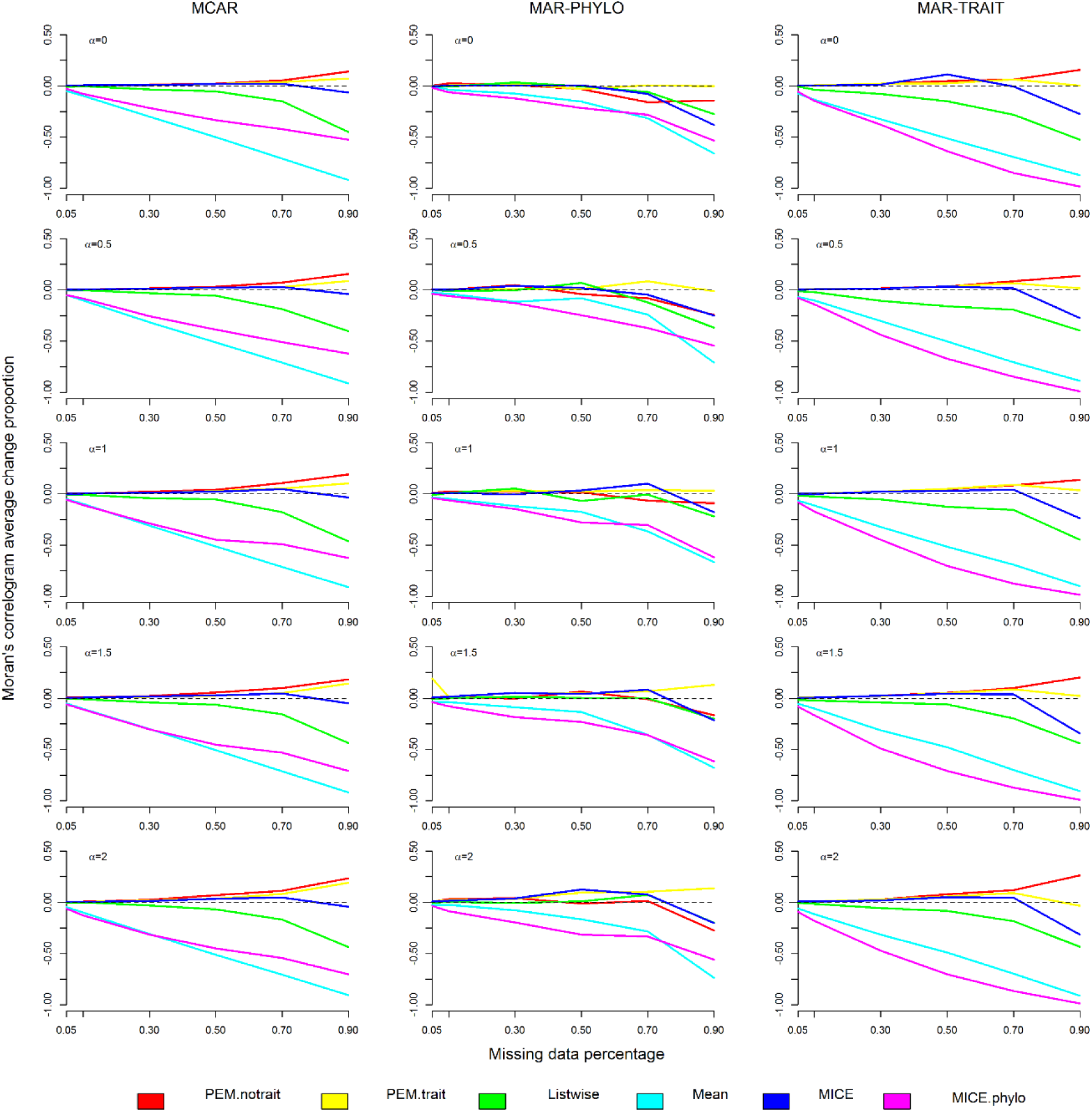
Moran’s Correlogram average change proportion under different methods, OU selective strength, missing data percentage and mechanisms.

**Figure 5.**
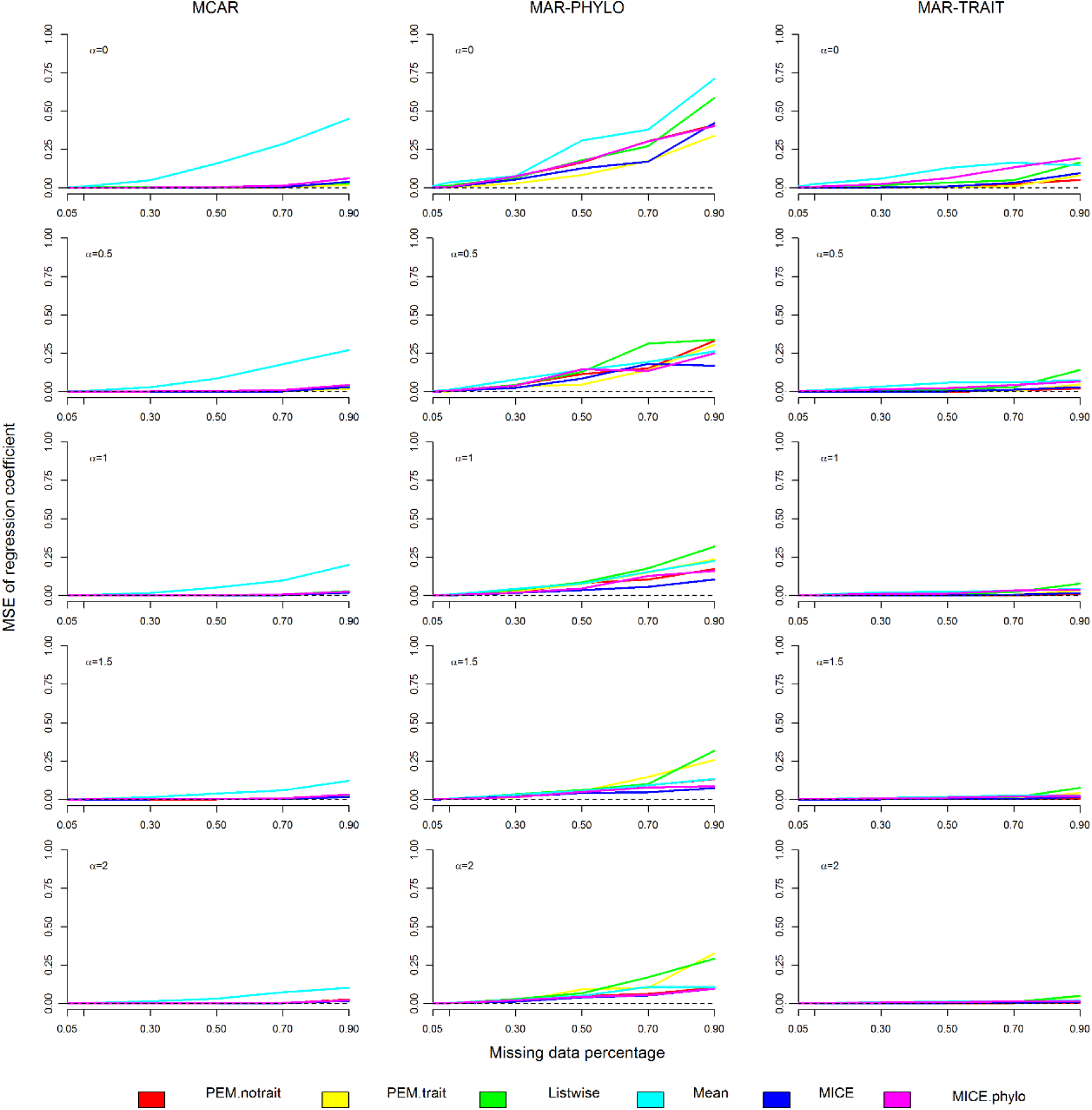
Regression coefficient MSE (mean squared error) under different methods, OU selective strength, missing data percentage and mechanisms.

The level of selection strength on trait evolution under the OU process was also important for the performance of imputation methods (Table 1). Accordingly, we found a tendency PS_delta_ and MSE to decrease as the selection strength increased from pure evolutionary drift (i.e. OU alpha = 0; Brownian motion) to strong selection (OU alpha = 2) (Fig. 3-5; Fig. S2 and S3, Appendix S2).

**Table 1.** Model selection of descriptive statistic and phylogenetic signal errors. The values represent the ∆AICc of the three best models for each error.

| Models | Blomberg's K | PS Moran | Imputation error | Mean | Variance | Regression Coefficient |
| --- | --- | --- | --- | --- | --- | --- |
| Model 1 | 6.23 | - | 0.00 | 0.04 | 11.71 | - |
| Model 2 | 0.00 | - | 4.71 | 0.00 | 0.00 | 8.28 |
| Model 3 | 3.79 | - | - | - | 1.94 | - |
| Model 4 | - | 0.00 | - | 15.45 | - | 0.00 |
| Model 5 | - | 1.07 | - | - | - | 3.74 |
| Model 6 | - | - | 34.85 | - | - | - |
| Model 7 | - | 3.31 | - | - | - | - |

The less sensitive methods were those that considered phylogenetic information in the imputation process (Fig. 3-5). PEM.trait, PEM.notrait, and MICE.phylo showed results less sensitive over different mechanisms of missing data (Fig. 3-5; Fig. S2 and S3, Appendix S2). From these three methods, PEM.trait was the less sensitive. The MEAN method was the most sensitive, similarly to MICE under MAR.PHYLO scenario (Fig. 3-5; Fig. S2 and S3, Appendix S2). The LISTWISE method caused the lowest changes in Blomberg’s K under all missing data mechanisms (Fig. 3) and for descriptive statistics only under the MCAR mechanism (Fig. 5; Fig. S2 and S3, Appendix S2).

Phylogenetic signal metrics (Blomberg’s K and PS_Moran_) were lower than the original (before imputation) when using MEAN and MICE methods (Fig. 3 and 4). All other methods estimated PS_Moran_ correctly under most simulated scenarios (Fig. 4), whereas the estimation of Blomberg’s K showed different patterns (Fig. 3). Blomberg’s K was overestimated by PEM.trait and PEM.notrait and underestimated by MICE, even under the MCAR missing mechanism (Fig. 3). Nevertheless, Blomberg’s K estimation errors decreased when phylogenetic eigenvectors were used in MICE.phylo (Fig. 3).

Descriptive statistics (mean, variance, and regression coefficient) were well estimated by all imputation methods (except MEAN) under MCAR. MAR.TRAIT and MAR-PHYLO generated biased estimations, but these biases were higher under MAR- PHYLO (Fig. 5, Fig. S2 and S3, Appendix S2). Nonetheless, variance had high estimations errors in MAR-PHYLO and MAR-TRAIT, independent of the imputation methods (Fig. S3, Appendix S2). For all descriptive statistics, considering phylogenetic structure improved estimations in MAR.PHYLO (Fig. S5, Fig. 2 and 3, Appendix S2).

Imputation error was lower when correlation (*r*) between the target and auxiliary traits was 0.9 than *r* equal 0.6. When traits were moderately correlated (*r* = 0.6), the lowest imputation error was found under missing completely at random (MCAR) scenarios (Fig. S4, Appendix S2) and when using imputation methods that considered phylogenetic information (PEM.trait, PEM.notrait, and MICE.phylo) (Fig. S4, Appendix S2). Moreover, all imputation methods performed better under the MAR.TRAIT than under MAR.PHYLO missing mechanism, but still poorly than under MCAR (Fig. S4, Appendix S2). When traits were strongly correlated (*r* = 0.9), the MICE methods presented lower imputation errors than when these traits were correlated moderately (*r* = 0.6). The PEM.notrait method increased its imputation errors when trait correlation was strong (*r* = 0.9) and PEM.trait was not influenced by correlation strength.

We found that estimation errors of descriptive statistics (MSE), Blomberg’s K, PS_Moran_ and imputation error (NRMSE) were influenced by all factors individually and their interactions (Table 1). Despite some differences among selected models in respect to the two- and three-way interactions, all models had interactions among all factors in some level (Table 2). Finally, we found that imputation errors were correlated with estimation errors (PS_delta_ and MSE) (Fig. 6). In addition, the imputation error and estimation error relationship evaluated here was asymptotic in log-scale, thus as imputation error increases the estimation error increases faster (Fig. 7).

**Table 2.**
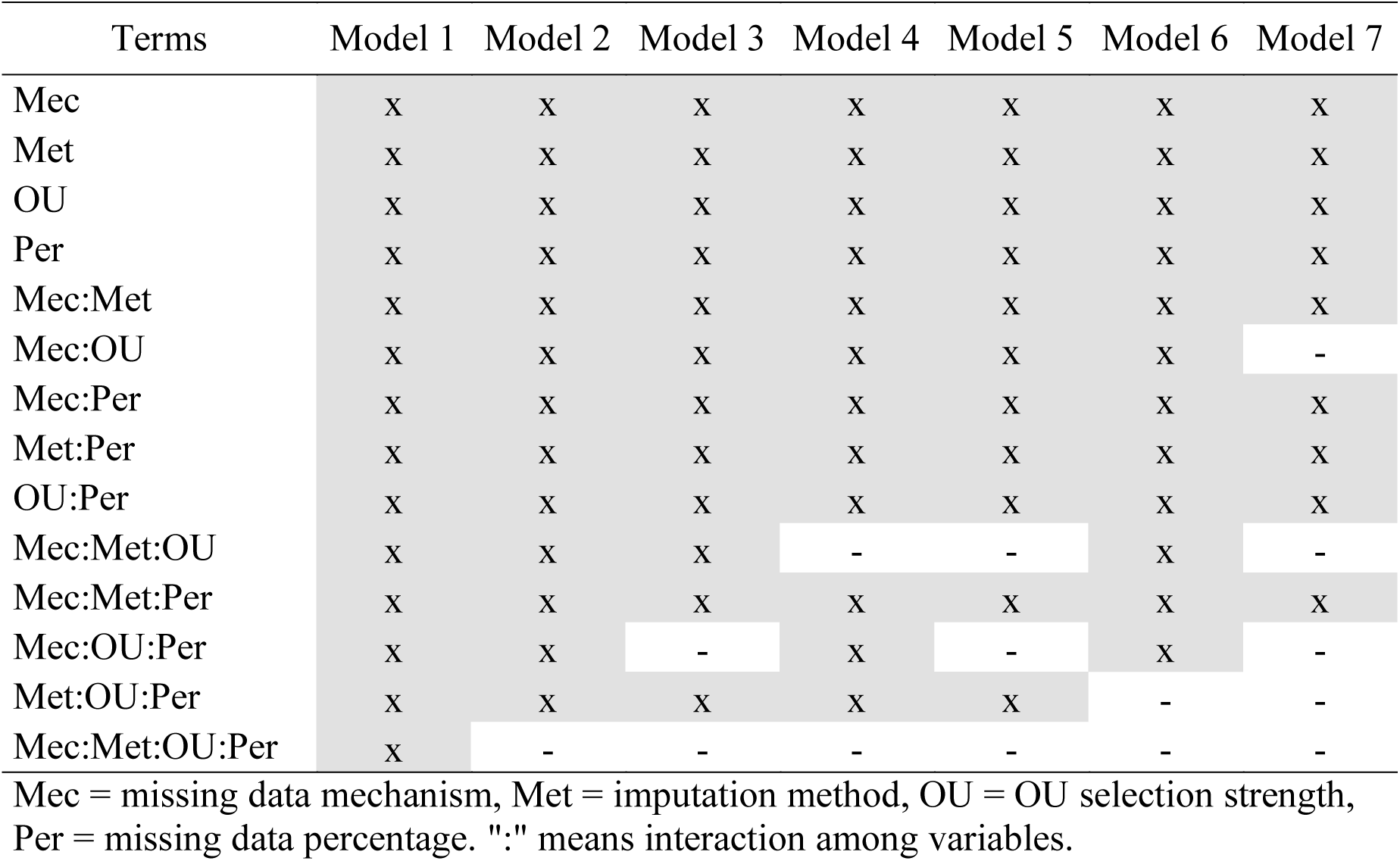
Description of selected models explaining estimation error of descriptive statistics (mean, variance and regression coefficient) and phylogenetic signal (Blomberg’s K and Moran Correlogram).

**Figure 6.**
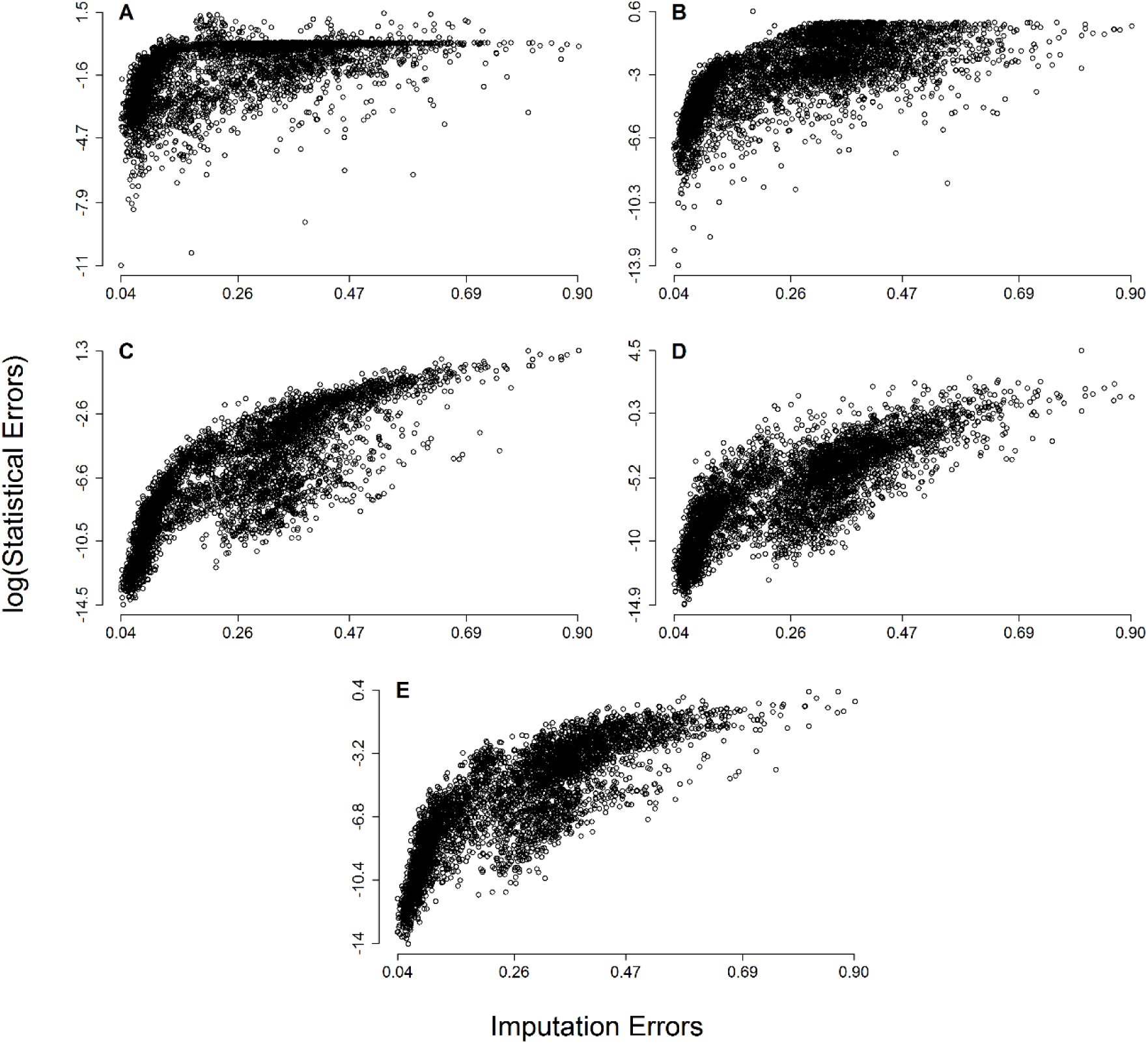
Scatterplot of imputation errors (average NRMSE) and statistical errors. (A) Logarithm of absolute average Blomberg’s K change proportion, (B) Logarithm of absolute average Moran’s Correlogram change proportion, (C) Logarithm of mean MSE, (D) Logarithm of variance MSE and (E) Logarithm of regression coefficient MSE.

**Figure 7.**
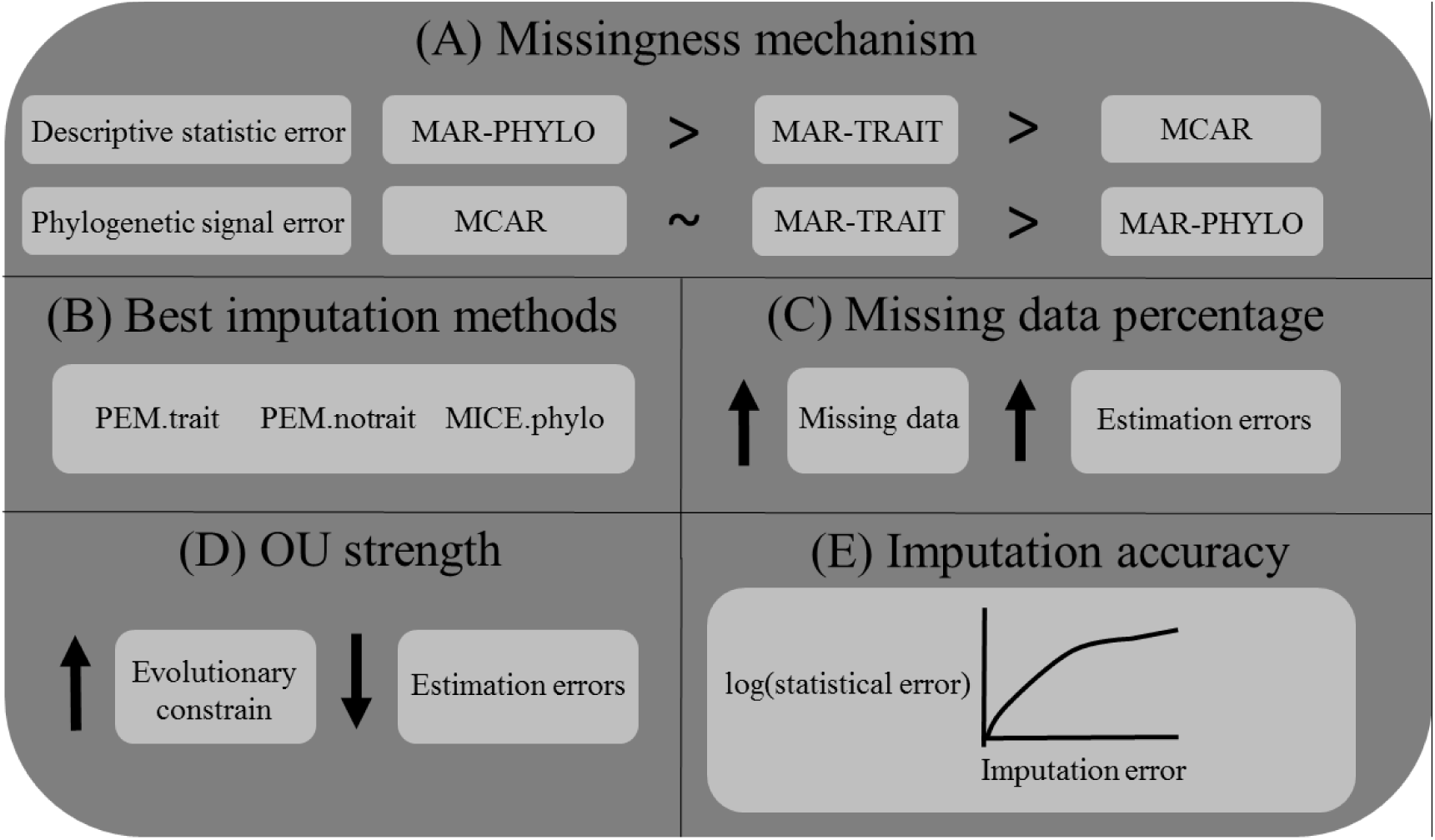

Summary of the main results showing (A) the differences on estimation errors among missing data mechanisms and estimated statistics; (B) highlighting the best imputation methods; (C) the effect of missing data percentage in statistical estimation; (D) OU selection strength; and (E) the non-linear relationship between imputation error and statistical estimation error logarithm.

## Discussion

Ecologists and evolutionary biologists are increasingly creating, using, and sharing large trait databases that are inevitably sparse and often completed by imputing missing values (Guénard *et al.* 2013; Swenson 2014; Schrodt *et al.* 2015). Here we argue that we should be extremely careful when using imputed databases, even for the estimation of simple parameters (i.e. means, variances and regression coefficients). Our findings revealed that estimations based on imputed data depends on every aspect of data property and strategy of analysis, as percentage of missing data, source/mechanism of absence, trait evolution, methods for gap filling, and statistics or parameters to be estimated. This has commonly been acknowledged in statistical research (Rubin 1976; Enders 2010) and should begin to be so in the ecological and evolutionary research as claimed by Nakagawa & Freckleton (2008). Based on our results, we can infer that the large changes in the estimations, due to different analytical choices, may also be an important cause of irreproducibility in our field (Borregaard & Hart 2016).

The most pervasive obstacle for deriving conclusions from large datasets is simply the proportion of those species lacking data. Previous studies found that reliable estimations from imputed data can be made when up to 60% of the values were missing (Barzi 2004; Penone *et al.* 2014). However, in our results, the effect of missing data percentage was not direct, but rather interacted with all of the other aspects evaluated here. Thus, there is no simple way of deriving a threshold on how much missing data would be allowed to be imputed and still make reliable estimations.

Knowing the causes of data absence is the first issue to be sorted out before any analysis (van Buuren 2012). The most common assumption in ecological and evolutionary studies is that data is missing completely at random (MCAR). This is evident in the wide variety of functions of the most commonly used software (the R programming language) allowing deleting missing values indiscriminately. Indeed, if data were under MCAR, previous findings and ours showed that estimations based on deletions and imputations could safely be made (Nakagawa & Freckleton 2010; Penone *et al.* 2014; Taugourdeau *et al.* 2014). However, biological data are rarely missing completely at random (Nakagawa & Freckleton 2008; Enders 2010). For instance, bias in ecological data absence can be related to the fact that some taxa are most studied than others (Gonzalez-Suarez *et al.* 2012). Moreover, such bias can stem from body mass differences among species, where large species have a higher probability of being described first (Vilela *et al.* 2014) and have their data collected (Gonzalez-Suarez *et al.* 2012) compared to small species. Also, species present in easily accessible regions are better studied than those occurring in regions that are hard to access (Reddy & Dávalos 2003). In our simulations, higher biased estimates were found when data were missing at random but correlated with other variable (MAR), especially phylogeny (MAR.PHYLO). Such results differ from those found by Penone *et al.* (2014), who did not find significant estimation differences among missing data mechanisms. This discrepancy could be related to our way of simulating MAR.PHYLO, creating a stronger phylogenetic structure than that simulated by them.

Our simulations revealed that imputation methods considering phylogenetic structure (PEM.trait, PEM.notrait and MICE.phylo) performed better than methods not doing so (MEAN, LISTWISE, and MICE) under all missing data mechanisms (MCAR, MAR.PHYLO, and MAR.TRAIT). Such findings support previous claims favoring “phylogenetic imputation” as a powerful tool in predicting missing species values (Penone *et al.*, 2014; Swenson 2014). More interestingly, our results showed that some phylogenetic imputation methods (PEM.notrait) perform better than non-phylogenetic ones, even when missing data was uncorrelated with phylogeny but to an auxiliary trait (MAR.TRAIT). This result was unexpected based on missing data theory, which suggests that under MAR.TRAIT some variable correlated with missing data probability is required to guarantee reliable estimations (Enders 2010).

Overall, PEM.trait performed best among all imputation methods tested. A potential caveat of this method is the imputation of a single value for each missing datum, thus not accounting for uncertainty of the imputed value. Consequently, PEM.trait (or PEM in general) may underestimate standard errors and bias subsequent hypothesis testing (i.e. increasing Type I error rates) (Enders 2010; van Buuren 2012). To avoid such biases, the statistical literature suggests using multiple imputation methods (Schafer & Graham 2002; Enders 2010; van Buuren 2012). However, our results did not show better performance of MICE, even when including phylogenetic information, in estimating descriptive statistics or phylogenetic signal compared to PEM. Despite multiple imputation being one of the most suggested methods for handling missing data (van Buuren 2012), additional research is necessary to evaluate its performance with phylogenetically structured data.

Filling missing values by averaging the observed ones (MEAN) or simply deleting species with missing values (LISTWISE) generated poor estimates, which is related to the fact that both methods assume that data is MCAR. MEAN only worked satisfactorily for estimating the trait average. LISTWISE disrupts the distribution of trait values, thus results in biased estimates (Enders 2010). However, this method performed well when estimating phylogenetic signal. This is encouraging, given that researchers interested in trait phylogenetic signal usually delete missing values (Blomberg & Garland 2002; Kamilar *et al.* 2013) thus guaranteeing potentially unbiased results.

Phylogenetic imputation is based on the assumption of target traits being phylogenetically structured (i.e. showing phylogenetic signal; Swenson 2014). However, phylogenetic structure is dependent on how traits evolved (Diniz-Filho 2001; Guénard *et al.* 2013). Accordingly, trait evolution was an important issue in our study. Across our simulated scenarios, estimation errors were higher when target traits were simulated under Brownian motion (BM) than under OU processes, agreeing with previous study (Guénard *et al.* 2013). Better estimates under OU than BM processes may result from higher trait resemblance and lower variance among species generated when increasing selection strength under OU processes (Hansen 1997; Butler & King 2004). Thus, predicting missing values of target traits will benefit from knowing their particular evolutionary model and will be more accurate if such traits evolved under strong selection regimes. Again, this suggests that researchers need to find the appropriate evolutionary model for their target traits before judging the need to use phylogenetic imputation methods for handling missing data. It should be noted, however, that fitting evolutionary models over incomplete data could itself be biased owing to the use of observed values only and thus pruned phylogenies (Slater *et al.* 2012).

Despite we showed phylogenetic imputation may recover descriptive statistics, phylogenetic imputation methods may produce bias when estimating phylogenetic signal. More specifically, our findings suggest that such methods can actually alter the original phylogenetic structure of the trait (i.e. the structure if data were complete). In fact, PS may be incorrectly estimated even under MCAR. Moreover, when using Blomberg’s K, imputation by PEM overestimated the original phylogenetic signal of the target trait (i.e. created when the trait was simulated) whereas MICE.phylo underestimated it.

In addition, PS estimation errors were dependent on the evaluated metric. Regardless of the simulated scenario, estimation errors were lower for PS based on Moran’s I correlogram than Blomberg’s K. Similarly, Münkemüller *et al.* (2012) showed that Moran’s I is less sensible than Blomberg’s K to changes in trait phylogenetic structure even when random noise is added. Blomberg’s K measures a global pattern along a phylogeny, based on observed and expected total trait variance under Brownian motion (Blomberg *et al.* 2003), whereas Moran’s I correlogram measures the correlation of trait values within different phylogenetic distance classes (Gittleman & Kot 1990). Therefore, changes in total trait variance caused by imputation may not have strong impacts on within-class correlations, rendering Blomberg’s K more sensitive than Moran’s I to such changes.

New proposed methods to fill sparse databases currently concerns about their degree of imputation error, that is how much imputed values deviate from the original trait values (Guénard *et al.* 2013; Penone *et al.* 2014; Schrodt *et al.* 2015). We found that single and multiple phylogenetic imputation methods can be highly accurate, resulting in small deviations between imputed and observed values, as suggested by other authors (Guénard *et al.* 2013; Penone *et al.* 2014; Diniz-Filho *et al.* 2015; Schrodt *et al.* 2015). In addition, we found that imputation error was positively correlated with estimation errors but their relationship was not linear. That is, increasing imputation error causes estimation errors to increase much more rapidly. This is particularly relevant if researchers were to use imputed databases blindly – without correctly treating imputed values. Such practice could create spurious results. This is because even if imputation is accurate, imputed values simply represent one among several possibilities without providing information on imputation uncertainty. In fact, using an accurately imputed database does not necessarily mean that the original trait distribution and its relationship with other variables will be recovered (van Buuren 2012).

## Concluding remarks

Instead of providing imputed trait databases, we should focus on treating missing values with appropriate methods. We have shown here that such methods should consider phylogenetic information. With the increase of computational literacy among ecologists and evolutionary biologists (Ram 2013), we encourage researchers to use simulations of their data and methods to find the appropriate solution for their study goals. Furthermore, researchers need to develop phylogenetic methods that consider imputation uncertainty and preserve the original data’s phylogenetic signal. Missing data is one of the most pervasive features of trait databases and the only effective solution for this Raunkiaeran shortfall is collecting more data. Nevertheless, acknowledging such shortfall instead of ignoring it will effectively help guiding research towards solving it.

## Acknowledgements

We thank G. Guénard for advice on PEM and L. Revell for help on evolutionary simulations. L.J. was supported by a CAPES doctoral fellowship. FV was supported by a BJT “Science without borders” CNPq grant. L.M.B. and J.A.F.D.F. are continuously supported by CNPq productivity grants.

